# Antibody Mediated Diversification of Primary and Secondary Humoral Immune Responses

**DOI:** 10.64898/2025.12.15.694384

**Authors:** Dennis Schaefer-Babajew, Laurine Binet, Gabriela S. Silva Santos, Chiara Ruprecht, Lachlan P. Deimel, Mohamed A. ElTanbouly, Dounia Gharrassi, Gabriella Lima dos Reis, Clara Uhe, Kai-Hui Yao, Brianna Hernandez, Parul Agrawal, Anna Gazumyan, Leonidas Stamatatos, Harald Hartweger, Michel C. Nussenzweig

## Abstract

Humoral immune responses are characterized by increasing antibody affinity and diversity over time. Increased affinity is mediated by a combination of immunoglobulin gene somatic mutation and iterative cycles of selection in germinal centers. Less is understood about how diversity increases. Here we examine the role of antibody feedback in diversifying immune responses in mice that produce B cells that are incapable of secreting antibodies. To this end, we produced two strains of mice, one that expresses only membrane and secreted forms of IgM, and a second that produces only the membrane bound form of IgM. Analysis of primary and secondary immune responses show that antibody feedback significantly diversifies both primary and secondary immune responses even when antibodies are present at levels that are 10–30 fold lower than physiologic. The data have significant implication for sequential vaccination approaches aimed at shepherding immunity to produce broadly neutralizing antibodies to highly diversified pathogens such as HIV-1 and Influenza.

**Summary:** Humoral immune responses diversify over time but whether secreted antibodies influence this process is unknown. Using antibody secretion-deficient mice this study shows a profound impact of secreted antibodies on the evolution of B cell diversity after vaccination.

## Introduction

Both the affinity and diversity of antibodies elicited by immunization or infection increase over time by a process that involves iterative cycles of selection, clonal expansion and somatic mutation in germinal centers (GCs) (Bannard and Cyster, 2017; Victora and Nussenzweig, 2022). These two apparently discordant features of humoral immune responses, increasing affinity and diversity, can be accounted for by dynamic changes in selection over time and by the existence of two cellular compartments, plasma cells and memory B cells respectively. Plasma cells develop by a mechanism that ensures increasing circulating antibody affinity (Zotos and Tarlinton, 2012; Nutt et al., 2015; Kräutler et al., 2017; Ise et al., 2018; Tas et al., 2022; ElTanbouly et al., 2024; MacLean et al., 2025). Memory B cells are selected by a separate mechanism that emphasizes diversity over affinity (Inoue et al., 2022, 2021; Suan et al., 2017; Viant et al., 2020; Wang et al., 2017).

Antibody diversification becomes increasingly important during secondary or booster immune responses, when only small numbers of high affinity antigen binding cells can be detected in GCs (Dal Porto et al., 2002; Eckl-Dorna et al., 2019; Mesin et al., 2019; Malladi et al., 2025). Given the strong selection pressure imposed by iterative cycles of mutation and cell division in the GC, the evolution away from high affinity is counterintuitive. However, diversification may be an important evolutionary adaptation to deal with rapidly changing pathogens such as Coronaviruses and Influenza virus (Heyman, 2000; Cyster and Wilson, 2024).

Several mechanisms have been suggested to account for the increase in diversity and loss of detectable antigen binding in late primary and secondary GCs. These include: 1. Continual recruitment of additional lower affinity cells during the reaction (De Carvalho et al., 2023; Hägglöf et al., 2023); 2. Increasing T cell fitness that lowers the threshold of B cell selection (Woodruff et al., 2018; Merkenschlager et al., 2023); 3. Antibody mediated antigen masking or enhancement (Heyman, 2000; Cyster and Wilson, 2024).

The effects of antibody feedback on immune responses have been studied since 1909 when they were first described by Theobald Smith in experiments on Diphtheria toxin (Smith, 1909). Recent data shows that masking appears to be mediated by high affinity antibodies and enhancement by low concentrations or low affinity antibodies that may be present even before immunization or in the early stages of the immune response (Heyman, 2000; Cyster and Wilson, 2024). But because all experiments to date, including Smith’s, have been performed in organisms that have at least some level of circulating antibody, precisely how masking and enhancement contribute to the evolution of humoral immunity and GC dynamics remains to be determined. Understanding these phenomena is increasingly important in developing strategies for sequential vaccination for difficult pathogens such as malaria parasites, HIV-1 and influenza virus where antibody mediated masking can interfere with the evolution of immunity in response to serial vaccine boosters (Cyster and Wilson, 2024).

Here we examine the role of antibody feedback in regulating GC responses in mice that have an intact polyclonal B cell compartment but do not secrete antibodies.

## Results

### Antibody secretion-deficient mice

To examine how secreted antibodies impact the development of immune responses, we engineered the IgH locus to produce two strains of mice: a control strain that expresses membrane and secreted forms of IgM (M-only) but no other isotypes; and a second strain whose B cells express only the membrane-bound form of IgM (mM-only) (Fig. 1 A). To this end, the C57BL/6J Ig heavy chain locus was modified using CRISPR-Cas9 to delete a 150 kb region 5’ of *Ighd* exon 1 to the 3’ UTR of *Igha*, leaving only the IgM locus intact (M-only mice, see methods for further details). M-only mice were subsequently engineered to produce the mM-only mice by removing the stop codon of the secreted splice form and its intronic polyadenylation signal in *Ighm* exon 4 (Danner and Leder, 1985; Nussenzweig et al., 1987; Boes et al., 1998; Ehrenstein et al., 1998; Waisman et al., 2007). The later abrogates production of the secreted form by preventing its termination thereby enforcing splicing to produce the membrane-bound isoform of IgM. We focused on the IgM isotype because it does not bind to most Fc receptors and therefore its Fc-mediated activity is generally more limited than other isotypes.

**Figure 1.**
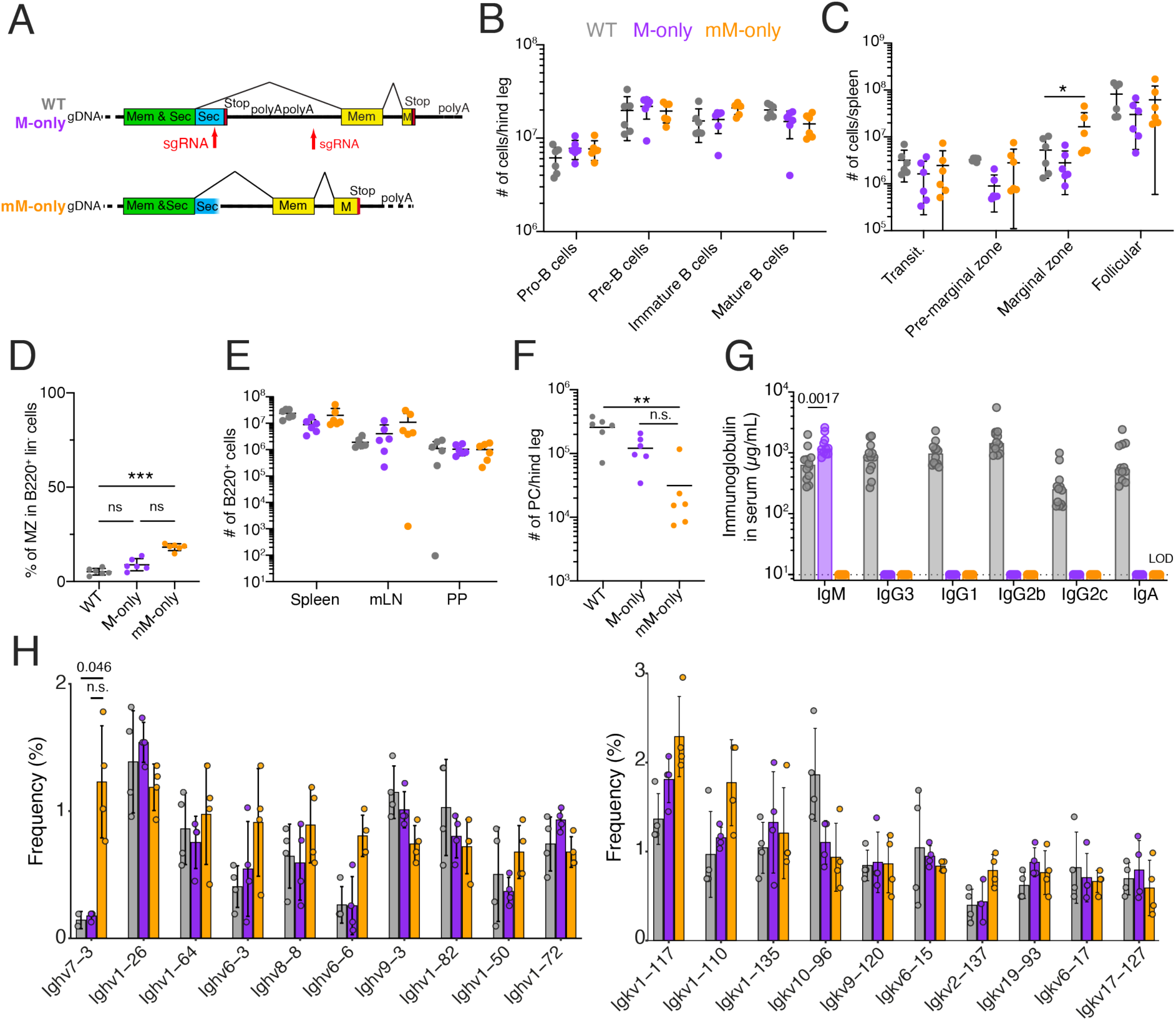
B cell development in M-only and mM-only mice. (**A**) Schematic shows structure of *Ighm* exon 4 in WT, M-only and mM-only mice. (**B**) Number of pro-, pre-, immature and mature B cells in the bone marrow on one leg by flow cytometry. Gating as in Fig. S1 A. (**C**) Number of the splenic B cell subsets by flow cytometry. Gating as in Fig. S1 B. (**D**) Percentage of splenic marginal zone B cells. Gating in Fig. S1B. (**E**) Number of live B cells in spleen, mesenteric lymph node (mLN) and Peyer’s Patches (PP) by flow cytometry. Gating in Fig. S1B-D. (**F**) Number of plasma cells in the bone marrow of one leg by flow cytometry. (**G**) ELISA quantification of total IgM, IgG3, IgG1, IgG2b, IgG2c and IgA in the serum of WT, M-only and mM-only mice. (**H**) Bar graph depicting the relative abundance of Ighv (left panel) and Igkv (right panel) gene usage in follicular B cells. Top ten most frequent genes ranked by the mean frequency in mM-only mice are shown. Data was pooled from two independent experiments. Each dot or circle represents a mouse. P-values of non-parametric one-way ANOVA with Tukey’s Honest Significance test are shown. All experiments were performed at least twice. **p <0.01 ***p<0.001. Bars indicate mean ± standard deviation (SD).

Consistent with the observation that membrane-bound IgM regulates B cell development in the bone marrow (Nussenzweig et al., 1987; Kitamura et al., 1991), M-only and mM-only mice showed generally normal numbers of pro-B, pre-B, immature and mature B cells in the bone marrow (Fig. 1 B and Fig. S1 A). Moreover, the overall numbers of transitional and mature B2 cells in spleen were similar, although a minor decrease of T2 cells was observed in mM-only mice. As reported by others, mice lacking secreted Igs, show increases in splenic B1 and marginal zone B cells but total numbers of B cells were similar in spleen, mesenteric lymph nodes (mLN) and Peyer’s patches. (Fig. 1 C-E and Fig. S1 B-G) (Hannum et al., 2000; Anderson et al., 2006). *κ*/*λ* light chain ratios showed minor differences in mM-only mice compared to both WT and M-only mice (Fig. S1 H). There were fewer plasma cells in the bone marrow of M-only mice compared to WT and they were difficult to detect in mM-only mice in (Fig. 1 F, Fig. S1 A). Serum ELISAs performed under steady state conditions showed slightly increased IgM but no other isotypes in M-only mice and no detectable antibody in mM-only mice (Fig. 1 G, P=0.0017). Despite the absence of other isotypes, single-cell Ig mRNA sequencing did not reveal any notable differences in Ighv, Igkv and Iglv usage by follicular B cells between wild type (WT), M-only and mM-only mice except for an enrichment of Ighv7-3 in mM-only mice compared to WT (Fig. 1 H and Fig. S1 I). mM-only B cells showed higher levels of surface IgM expression in both bone marrow and the spleen possibly due to their lack of IgD and inability to secrete (Fig. S1 J). Finally, at steady state, mLN GCs in mM-and M-only mice had similar dark zone (DZ)/light zone (LZ) ratios and number of T follicular helper (T_FH_) cells (Fig. S1 K and L). In summary, B cell development appears largely normal, but there are no serum antibodies and few if any plasma cells in mM-only mice.

### Primary immune responses

To determine how antibodies impact the development of primary B cell immune responses we immunized mice with the Wuhan-Hu-1 SARS-Cov2 receptor binding domain (RBD) (Fig. 2 A). Antigen-specific antibodies were measured by ELISA (Fig. 2 B-D). As expected, M-only mice were limited to the IgM isotype, and mM-only mice produced no serum antibodies in response to the immunization (Fig. 2 B-D). The kinetics of the serological response in M-only mice was similar to WT (Fig. 2 B and D), but the total amount of antibody was lower when considering all isotypes (Fig. 2 B). Notably, whereas the total antibody levels in WT mice decreased over the 150-day period of observation, serum Ig levels were relatively stable and even increased modestly in M-only mice (Fig. 2 B).

**Figure 2.**
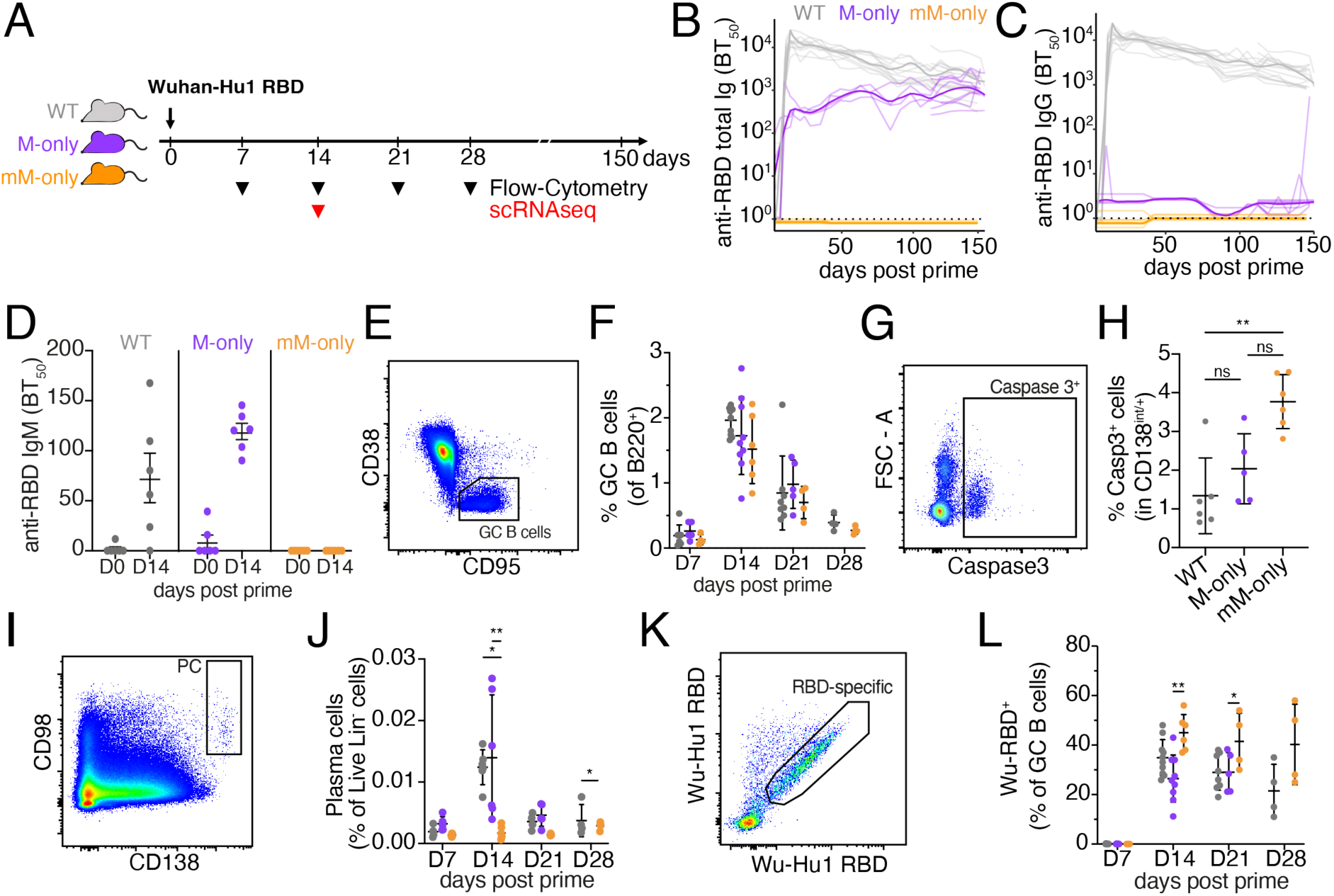
Primary vaccine responses. (**A**) Diagram of the experimental protocol. (**B**) ELISA quantification of anti-RBD antibodies in the serum measured every 3 to 5 days between D0 and D150. (**C**) As in (B) for IgG anti-RBD antibodies in the serum. Thin lines (in B and C) represent single mice, bold lines represent group mean. (**D**) Serum anti-RBD IgM ELISA 0 or 14 after immunization. (**E**) Flow cytometric gating for CD38^−^CD95^+^ GC B cells. Pre-gated on B220^+^ cells in draining lymph nodes (dLN). (**F**) Quantification of GC B cells from (E) at the indicated time points after immunization. (**G**) Flow cytometry gating for Caspase3^+^ cells among CD138^int/+^ live Dump^−^ cells. (**H**) Quantification of (G) among CD138^int/+^ live Dump^−^ cells in dLN 14 days after immunization. (**I**) Flow cytometric gating for CD98^+^ CD138^+^ PC pre-gated on live Dump^−^ cells in the dLN. (**J**) Quantification of (I) at the indicated times after immunization. (**K**) Flow cytometric gating of RBD^+^ cells among GC B cells in the draining LNs. Pre-gated for GC as in (E). (**L**) Quantification of (K) at the indicated time points after immunization. Each dot in all dot plots represents one mouse. P-values of non-parametric one-way ANOVA are shown. All experiments were performed at least twice. Bars in (D, F, H, J and L) indicate mean ± SD. **p <0.01, *p<0.05.

Germinal center responses were analyzed by flow cytometry on days 0, 7, 14, 21 and 28 after immunization. GCs were absent from popliteal LNs before immunization (Fig S2 A). The absolute numbers of B cells and the relative proportions of germinal center B cells among B220^+^ B cells was similar among WT, M-only and mM-only mice at all time point points analyzed (Fig. 2 E and F and Fig. S2 A-C). Thus, the contribution of antibodies to GC magnitude and kinetics in the primary response to SARS-CoV2-RBD appears to be limited.

Plasma cell precursors were evident in GCs of all 3 strains, but showed increased caspase expression in mM-only mice and these mice had few detectable mature plasma cells in lymph nodes (LNs) or bone marrow (Fig. 2 G-J and Fig. 1 F and Fig. S2 D,E and Fig. S1 A). The data are consistent with the idea that plasma cells that are unable to secrete antibody die by apoptosis (Iwakoshi et al., 2003; Todd et al., 2009; Bonaud et al., 2023).

We analyzed the antigen-binding capacity of GC B cells in the primary response by flow cytometry using the Wuhan-Hu-1 RBD labeled with 2 different fluorophores (Fig. 2 K and L). At the peak of the GC response, 14 days after immunization, there were similar proportions of RBD-binding cells in WT and M-only mice (35 % and 26 % respectively), but significantly more in the GCs of mM-only mice (45 % p=0.0021, Fig. 2 L and Fig. S2 F). Moreover, whereas antigen binding B cells decreased over time in WT and M-only mice their relative proportion appeared to persist in mM-only GCs (Fig. 2 L). Differences in BCR expression did not account for this difference in antigen binding as IgM surface expression levels were equivalent among genotypes (Fig. S2 G and H). The data is consistent with the idea that secreted antibodies mask immunodominant epitopes and increase antigen valency, thus curbing the relative competitive advantage of high-affinity immunodominant antigen binding cells in the later stages of the primary GC reaction (Heyman, 2000; Andrews et al., 2015; Schaefer-Babajew et al., 2023; Cyster and Wilson, 2024; Yan et al., 2025).

To gain further insights into the effect of antibodies on primary GC responses we examined clonality, diversity and IgV gene usage by single cell VDJ sequencing 14 days after immunization. As expected, expanded clones of B cells were found in GCs of all 3 genotypes (Fig. 3 A-C and G and Fig. S2 I). However, the individual clones appeared to be larger and less diverse in the GCs of mM-only mice than in mice that expressed secreted antibodies as evidenced by significant higher overall clonality and a decrease in the fraction of individual sequences (Fig. 3 B-C). Consistent with this observation mM-only GC B cells showed reduced Shannon and Inverse Simpson diversity indices, but similar levels of somatic mutation compared to their counterparts, even when analyzing the sequences of clones composed from the top 10 most frequent V genes in mM-only mice (Fig. 3 D-F and Fig. S2 J). We did not detect significant skewing of IgV gene usage (Fig. 3 G and Fig. S2 K). In conclusion, secreted antibodies appear to be essential for B cell repertoire diversification in primary GC responses to SARS-CoV2 RBD – a mechanism that appears to be significant even at the early stages of the response.

**Figure 3.**
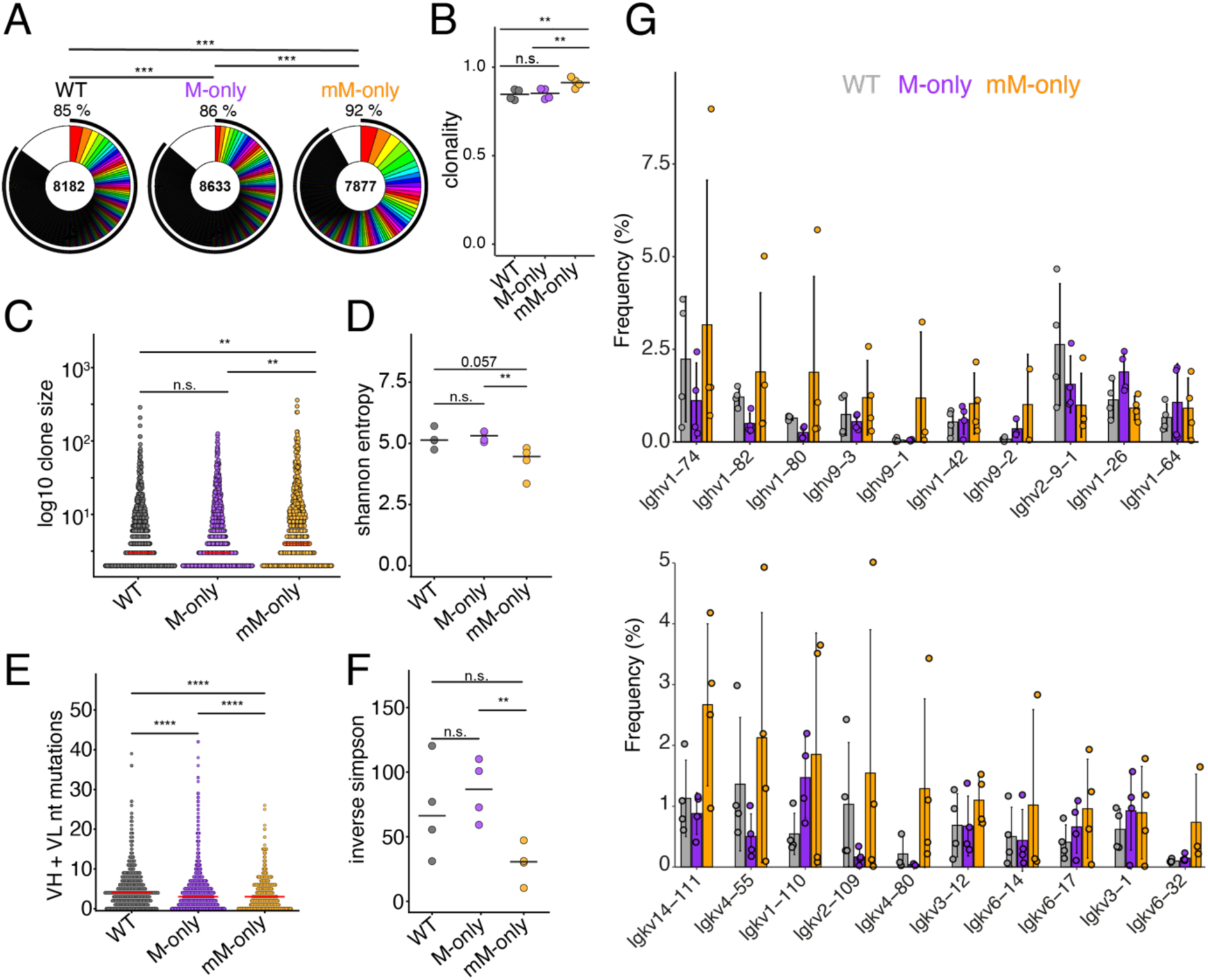
Primary GC antibody sequences. (**A**) Pie charts depicting the distribution of antibody sequences pooled from 4 mice/genotype obtained 14 days after prime immunization from GC B cells. Inner circle numbers indicate the number of sequences analyzed for each genotype. White section indicates sequences non-expanded sequences, colored or black pie slices are proportional to the number of clonally related sequences. The outlined black line indicates the percentages of cells in a clonal family. (**B**) Dot plot illustrating the percentage of clonally expanded sequences within each mouse. Each dots indicates one mouse. Bars indicate mean. (**C**) Dot plot showing clonotypes size distribution. Bars indicate median. Each dot indicates a clone. (**D**) Dot plot showing Shannon entropy scores for the sequences depicted in (A) within each mouse. Bars indicate median. (**E**) Dot plot showing combined VH + VL gene nucleotide mutations among sequences show in (A). Each dot indicates an antibody VH+VL pair. Bars indicate median. (**F**) Dot plot depicting inverse Simpson index for sequences shown in (A) for each mouse. Each dots indicates one mouse. Bars indicate median. (**G**) Bar graphs showing the relative abundance of mouse Ighv and Igkv gene usage among sequences shown in (A). Top ten most frequent genes ranked by the mean frequency in mM-only mice are shown. Bars indicate mean ± SD. Statistics in (A) indicate chi-squared test with Monte Carlo p-value simulation, p-values were subsequently corrected using the Benjamini-Hochberg (BH) procedure. Statistics in (B)-(F) indicate two-sided Mann-Whitney U test. ****p<0.0001, ***p <0.001,**p <0.01, *p<0.05.

### Secondary Immune Responses

To determine whether antibodies developing during the primary response influence subsequent booster responses, we vaccinated WT, M-only and mM-only mice in accordance with protocols used during human immunization against SARS-CoV-2 using full-length stabilized SARS-CoV-2 spike protein (Fig. 4 A).

**Figure 4.**
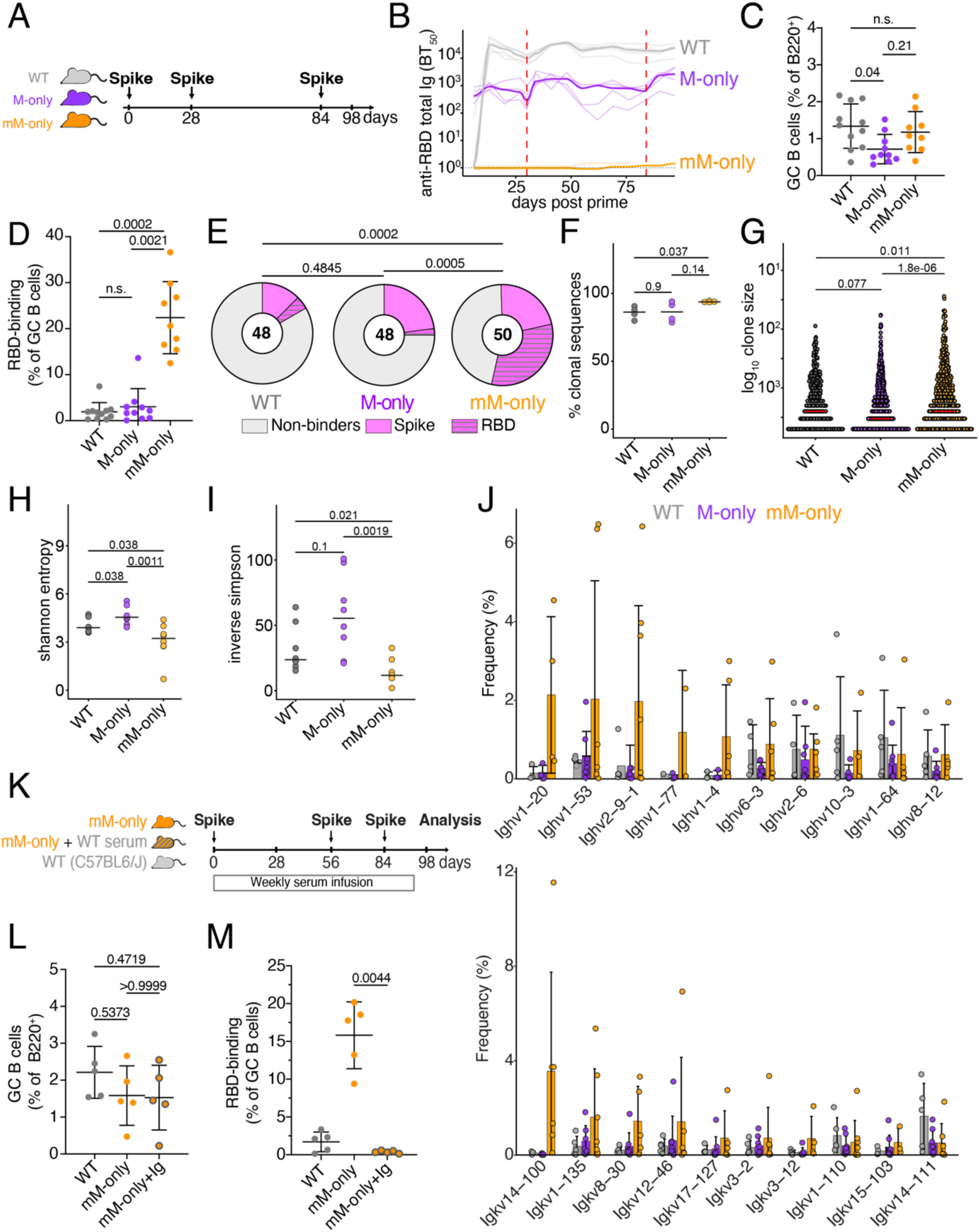
Prime boost vaccination. (**A**) Diagram of the experimental protocol for (A) to (J). (**B**) ELISA quantification of total anti-RBD antibodies in the serum measured weekly from D6 to D97 after prime immunization. Thin lines represent single mice, bold line indicates group mean. Dashed red lines indicate booster immunizations. (**C**) Percentage of GC B cells among B220^+^ B cells in the draining LN at D98. Gating as in (Fig. 2 E). (**D**) Percentage of RBD-binding GC B cells in the draining LN at D98. Each dot represents one mouse. Gating as in (Fig. 2 K). Each dot represents one mouse. Bars indicate mean ± SD (**E**) Pie charts depicting the distribution of RBD-binding, Spike-binding or non-binding recombinant antibodies cloned from the largest expanded clones of 4 mice/group of GC B cells on D98 from draining LNs. BT_50_ < 10 µg/mL considered binders. Inner circle numbers indicate the number of recombinant antibodies analyzed. (**F**) Dot plot showing the percentage of clonal sequences from GC B cells on Day 98. Each dot represents one mouse. Bars indicate median. (**G**) Number of mutations in GC B cells on D98. Each dot represents one clone. Bars indicate median. (**H**) Dot plot depicting the Shannon entropy for sequences from (F). (**I**) Dot plots depicting the inverse Simpson index for sequences from (F). Each dot represents one mouse. Bars indicate median. (**J**) Bar graphs showing the relative abundance of mouse Ighv and Igkv gene usage among sequences from (F). Top ten most frequent genes ranked by the mean frequency in mM-only mice are shown. Bars indicate mean ± SD. (**K**) Diagram of the experimental protocol for (L) and (M). (**L**) Percentage of GC B cells among B220^+^ B cells in the draining LN on day 98. Each dot represents one mouse. Gating as in (Fig. 2 E). Bars indicate mean ± SD (**M**) Percentage of RBD-binding GC B cells. Each dot represents one mouse. Gating as in (Fig. 2K). Bars indicate mean ± SD. Statistics in (C), (D), (J), (L) and (M) indicate non-parametric, one-way ANOVA p-values. Statistics in (E)-(I) indicate two-sided Mann-Whitney U test. ****p<0.0001, ***p <0.001,**p <0.01, *p<0.05. All flow cytometry experiments are from at least 2 independent experiments.

Serum antibody responses were measured by ELISA at approximately weekly intervals from days 6 to 97 after immunization. As in the primary responses, antibodies were not detectable in mM-only mice (Fig. 4 B and Fig. S3 A), and M-only mice produced approximately 30-fold less serum antibody compared to WT mice 14 days after boosting. Flow cytometry on cells obtained from draining lymph nodes showed that M-only had lower proportions of GC B cells, but mM-only was not significantly different from WT (Fig. 4 C). Thus, the complete absence of secreted antibodies does not impact the ability to form or sustain secondary GCs.

Antigen-binding GC B cells were enumerated using fluorescently labeled Wuhan-Hu-1 RBD since this is the primary target of the anti-Spike immune response in mice and humans. Notably, the 3 strains differed significantly in the fraction of Wuhan-Hu-1 RBD-binding B cells (Fig.4 D). Whereas only 1.9 % and 3 % of the B cells in WT or M-only GCs bound to RBD, 22 % of mM-only GC B cells showed demonstrable binding (p= 0.002, Fig. 4 D).

To determine the antigen-binding specificity of GC B cells that develop after boosting, we cloned and expressed 146 representative antibodies and tested them for binding to Wuhan-Hu-1 spike and the isolated RBD by ELISA (Table S1). Altogether, only 17 % (wild-type) and 25 % (M-only) of mAbs isolated from secondary GCs exposed to endogenous antibodies showed measurable binding to the immunizing antigen, whereas 54 % of the mAbs isolated from mM-only GCs were of sufficiently high affinity to bind by ELISA (Fig. 4 E and Fig. S3 B). Whereas 32 % of antibodies obtained from mM-only mice bound to the immunodominant RBD, only 4.2 % and 2.1 % of mAbs from WT and M-only GCs did so, respectively (p=0.0002 and 0.0005 respectively, Fig. 4 E). The fraction of antibodies binding to non-RBD epitopes on the spike was 12.5 %, 23 % and 22 % in WT, M-only and mM-only mice, respectively. To examine the nature of the antibodies produced after booster immunization, we compared the Ig sequences from single GC B cells. GC B cells obtained from secondary GCs of mM-only mice were more clonal and significantly less diverse than those obtained from M-only mice (Fig. 4 F-I). Again, we did not detect significant differences in V gene usage and genotypes displayed similar numbers of somatic mutations (Fig 4 J and Fig. S3 C). The data suggest that antibody feedback inhibits accumulation of GC B cells that are restricted to the initial immunodominant epitope even when those antibodies are restricted to the IgM isotype and when total Ig concentrations are 30-fold lower than physiologic.

To determine whether passively transferred antibodies are sufficient to revert the mM-only phenotype, we repeated the prime boost experiment with Wuhan-Hu-1 spike and transferred contemporaneous serum from WT mice into mM-only recipients (Fig. 4 K). As determined by ELISA serologic reconstitution was only partial with an approximately 10-fold lower level of specific antibodies in the mM-only recipients compared to their WT counterparts (Fig. S4 A). Notably, while the fraction of GC B cells in draining LNs was similar between genotypes, passive antibody transfer completely reverted the mM-only phenotype resulting in near complete loss of Wuhan-Hu-1 RBD binding B cells in the mM-only GCs (Fig. 4 L and M and Fig. S4 B and C). We conclude that small amounts of polyclonal antigen-specific serum is sufficient to drastically change antigen binding specificity in the GC. This observation is consistent with the idea that secreted antibodies mask immunodominant epitopes and thereby diversify epitope usage in GCs during polyclonal immune responses.

Sequential immunization strategies designed to produce broadly neutralizing antibodies (bNAbs) to HIV-1 and influenza virus are currently being investigated in animal models and in the clinic. The multi-step vaccine concept being tested involves using mutated, high-affinity priming immunogens to recruit rare bNAb precursors followed by iterative administration of progressively more native antigens to shepherd B cell responses through a series of somatic mutations required to produce bNAbs (Escolano et al., 2016; Burton, 2019; Haynes et al., 2023). At the end of the sequence, antigen binding B cells should retain binding to the priming immunogen and in addition bind to all the different antigens used in the vaccine. This idea has yet to produce necessary broadly protective serologic responses in part because “off target” antibodies to alternative sites on the immunogen dominate the response after booster vaccination (Escolano et al., 2019, 2021). To determine how circulating antibodies might contribute to this effect, we immunized mice with a series of 4 HIV-1 CD4-binding site-targeting immunogens, the first of which is currently in early clinical testing (NCT05471076; 426c.DMRS.Core (TM4) antigen, Fig. 5 A) (McGuire et al., 2016). Immunizations were administered in the same anatomic location throughout and longitudinal ELISA analysis for antibodies binding to 426c.DMRS.Core showed that titers were present throughout the immunization protocol in WT and M-only but not mM-only mice (Fig. 5 B).

**Figure 5.**
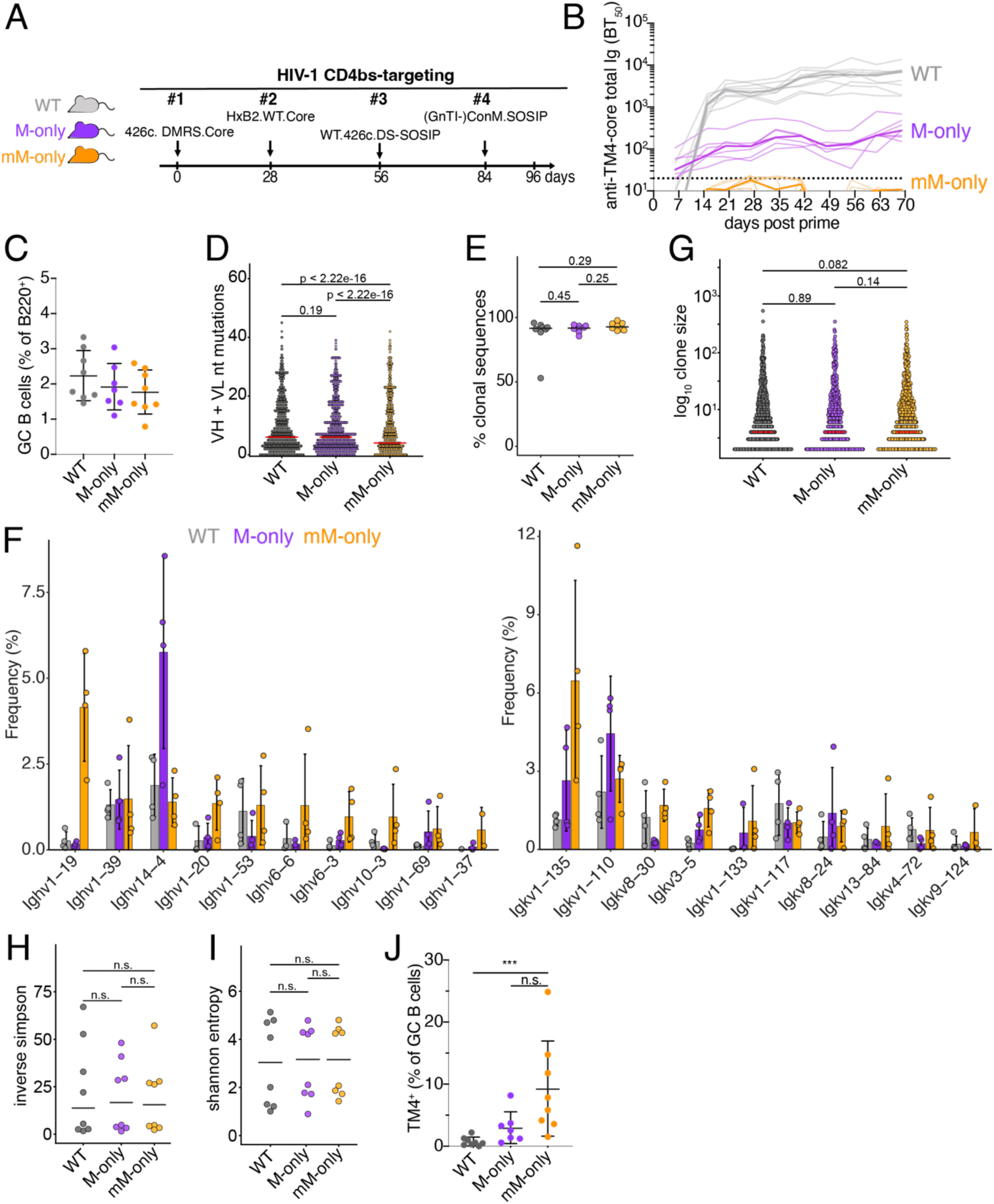
Sequential immunization with an HIV-1 antigen series. (**A**) Diagram of the experimental protocol. (**B**) ELISA quantification of total anti-TM4-core serum antibodies measured weekly from day 6 to 69. Thin lines represent single mice, bold lines represent group mean. (**C**) Percentage of GC B cells among B220^+^ B cells in the draining LN on day 96. Gating as in (Fig. 2 E). Bars indicate mean ± SD. (**D**) Dot plot showing combined VH + VL gene nucleotide mutations among GC B cells sequences on day 96. Each dot indicates an antibody VH+VL pair. Bars indicate median. (**E**) Proportions of clonal sequences within each mouse at day 96. Each dot indicates sequences from one mouse. Bars indicate median. (**F**) Bar graph showing the relative abundance of mouse Ighv left panel) and Igkv (right panel) gene usage in GC B cells on day 96. Top ten most frequent genes ranked by the mean frequency in mM-only mice are shown. Bars indicate mean ± SD (**G**) Dot plot depicting the percentage of clonal sequences from GC B cells on day 96. Each dot indicates a clone. Bars indicate median. (**H**) Dot plot depicting Inversed Simpson index of antibody sequences from day 96 GC B cells. Each dots represents one mouse. Bars indicate median. (**I**) Dot plot showing the Shannon entropy index form day 96 GC B cells. Each dots represents one mouse. Bars indicate median. (**J**) Percentages of TM4-binding GC B cells. p-values of non-parametric one-way ANOVA indicated. ***p <0.001.

14 days after the last boost GC B cell numbers were equivalent among the 3 mouse strains with median GC B cell frequencies between 1.7 % to 2.2 % of all B cells in the draining lymph node (Fig. 5 C and Fig. S5 A). Somatic hypermutation was again comparable across genotypes including expanded clones (Fig. 5 D and Fig. S5 B). Clonality, clone size, IgV gene usage and measures of diversity did not show significant differences (Fig. 5 E-I and Fig. S5 C), and considering all cells irrespective of HIV-1 antigen-binding, we saw a high degree of clonality in all 3 groups (Fig. 5 E). Antigen binding to TM4, but not to other sequentially delivered antigens was measured by flow cytometry. Notably, at the end of the sequential vaccine sequence WT, M-only and mM-only showed 0.76 %, 3 % and 9 % TM4-binding GC cells respectively (Fig. 5 J and Fig. S5 D). Thus, only mM-only mice that are devoid of circulating antibodies show continued levels of GC B cells that bind to the priming immunogen after a sequential series of HIV-1 vaccine boosters. In summary antibody feedback interfered strongly with persistence of antigen binding GC B cells in sequential immunization but differences in clonality and diversity were less pronounced compared to repeated SARS-CoV-2 spike protein immunization. We speculate that this difference arises partly from the nature of the antigens and reduced antibody feedback associated with the sequential use of different antigens. We conclude that circulating antibodies interfere with GC persistence of B cells that retain the ability to bind to the priming immunogen.

## Discussion

GCs are microanatomic compartments specialized for affinity maturation and diversification of humoral immune responses. Affinity maturation is an iterative process involving repetitive cycles of division and mutation in the GC dark zone followed by migration to the light zone where B cells test their newly acquired antigen receptors for binding and capture of antigen deposited in follicular dendritic cells. Selection is mediated by a combination of B cell receptor signaling and T follicular help (Victora and Nussenzweig, 2022; Chen et al., 2023). How GCs diversify antibody responses in the face of strong positive selection is less well understood. Our data indicate that even small amounts of specific polyclonal antibodies produced during the immune response are sufficient to diversify the GC. We hypothesize that this phenomenon is due to masking responses to immunodominant epitopes and by producing immune complexes that lower the affinity thresholds for B cell entry into the GC.

Antibody feedback has been studied primarily by passive antibody transfer in animals and humans (Heyman, 2000; Schaefer-Babajew et al., 2023; Cyster and Wilson, 2024). These experiments revealed that both IgM and IgG can produce immune complexes and mediate masking through interactions with Fc and complement receptors (Heyman, 2000). Moreover, recent work in antibody knock in mice producing antibodies of differing affinity to a selected epitope showed that the antibody feedback effects were dependent on affinity and that changes in the T cell compartment might also contribute (Barbulescu et al., 2025; Yan et al., 2025). In addition, even partial deletion of antibody secreting cells showed that masking impacts the composition of the GC (Schiepers et al., 2024). Our experiments extend these findings to primary and secondary immune responses in animals that have an otherwise intact polyclonal B and T cell compartment but are unable to produce secreted antibodies. Our data show that antibody feedback is a major mechanism for diversification of both primary and secondary humoral immune responses.

Diversification by antibody feedback has the potential to enable recall responses to rapidly evolving pathogens. For example, although initial responses to SARS-CoV-2 were focused on highly strain specific epitopes, antibody feedback diversified the response to include more conserved epitopes that offered some protection against newly arising variants. However, our data indicate that these feedback effects pose a very significant barrier to sequential immunization approaches such as those being tested for HIV-1 that aim to focus immunity to a singular broadly neutralizing epitope.

## Methods

### Mice

C57BL/6J mice were purchased from Jackson Laboratories. M-only and mM-only mice were created and maintained at the Rockefeller University with assistance from the Rockefeller University CRISPR and Genome Editing Center and Transgenic and Reproductive Technology Center. Cas9 (IDT) and sgRNA (IDT) ribonucleoprotein complexes targeting the IgH locus were electroporated into C57BL6/J single cell mouse embryos, which were then recovered and incubated overnight at 37°C before implantation into female foster mother mice. For M-only mice sgRNA spacer sequences GTAGATCTCTTCCTAAGAGG and TTACTAGGCTCCTCCATATG were used to delete *Ighd*–*Igha*. M-only mice were retargeted and with sgRNAs with spacer sequences CGCCTGTGTCAGACATGATC and GGGTAGGACAAGCAACGCAC to delete the part of *Ighm* exon 4 only found in the secreted IgM splice form, as well as the subsequent stop codon and alternative, intronic polyadenylation site (Danner and Leder, 1985; Boes *et al*., 1998; Ehrenstein *et al*., 1998; Waisman *et al*., 2007). Cutsite adjacent deletions were verified by PCR from tail DNA and Sanger sequencing of PCR products. Sequences were analyzed using Geneious Prime (GraphPad). Mice were bred to homozygosity. Presence or absence of antibody isotypes was verified by flow cytometry and ELISA of serum in homozygous animals from homozygous parents due to maternal transfer of antibodies in utero and through suckling.

Male and female mice aged 6-12 weeks were used. Animals were housed at the Rockefeller University Comparative Bioscience Center and all animal procedures were performed following protocols approved by the Rockefeller University Institutional Animal Care and Use Committee. Animals were housed at an ambient temperature of 22 °C and a humidity of 30–70 % under a 12 h–12 h light–dark cycle with free access to food and water. For experimental endpoints, mice were euthanized using CO_2_, followed by cervical dislocation.

### Immunization

All immunizations were performed by subcutaneous footpad injection of 5 µg of immunogen in 33 % alhydrogel (Invivogen).

### Serum infusion

Serum was obtained from immunized, time-matched C57BL/6J mice and pooled, before being administered to recipients at the equivalent time points. Each recipient received 200 μL of the pooled serum on a weekly basis.

### Flow cytometry

For steady state experiments, mice aged from 10 to 11 weeks were sacrificed and their spleen, mesenteric LN, Peyer’s patches and bone marrow were isolated and dissociated to obtain single-cell suspensions. Bone marrow from pelvic bone, femur and tibia was isolated by centrifugation at 10,000 g for 15 s. Spleen and Peyer’s patches were forced through a 70 µm cell strainer. LNs were collected into 1.5 mL Eppendorf tubes and dissociated using a pestle. Spleen and bone marrow pellets were then incubated in ACK lysis buffer for 15 min on ice, washed and processed together with the LN as described below.

For all experiments, single cell suspensions were incubated in Fc block (BD Biosciences) for 15 min. Cells were then centrifuged at 350 g for 5 min and resuspended and incubated in a solution of PBS diluted fluorescent-Biotin-antigen tetramers for 30 min on ice. Additional labeled antibodies were then added to the cells and samples were incubate for 30 additional minutes on ice. For intracellular staining of Caspase 3, cells were permeabilized using eBioscience Foxp3 permeabilization kit (reference: 00-5523-00), washed in the supplied permeabilization buffer and stained for 30 min at 4 °C. as per manufacturer’s instructions. The different cell populations were identified as follows: ProB (Dump^−^, CD19^+^, B220^+^, IgM^−^, CD2^−^), PreB (Dump^−^, CD19^+^, B220^+^, IgM^−^, CD2^+^), Immature B (Dump^−^, CD19^+^, B220^+^, IgM^+^, CD2^+^, CD23^−^), Mature B (Dump^−^, CD19^+^, B220^+^, IgM^+^, CD2^+^, CD23^+^), B1a (Dump^−^, CD19^+^, B220^−^, CD23^−^, CD5^+^), B1b (Dump^−^, CD19^+^, B220^−^, CD23^−^, CD5^−^), B2 (Dump^−^, CD19^+^, B220^+^), Transitional B (Dump^−^, CD19^+^, B220^+^, CD93^+^), pre-marginal zone B (Dump^−^, CD19^+^, B220^+^, CD21^hi^, CD93^−^, IgM^hi^, CD23^+^), marginal zone B (Dump^−^, CD19^+^, B220^+^, CD21^hi^, CD93^−^, IgM^hi^, CD23^−^), Follicular B (Dump^−^, CD19^+^, B220^+^, CD21^int^, CD93^−^, IgM^+^), PC at steady state (Dump^−^, CD98^+^, TACI^+^) or after immunization (Dump^−^CD98^+^ CD138^+^), GC B cells (Dump^−^, CD98^−^, CD138^−^, B220^+^, CD95^+^, CD38^−^). The Dump channel contains Ly6G, F4/80, NK1.1, CD4, CD8 antibodies and Zombie-NIR live/dead marker. Flow cytometric reagents are listed in Table S2.

Antigen tetramer staining was performed with a combination of two different fluorescently-labelled streptavidin conjugates per antigen. Biotinylated antigens at 5 µg/mL were individually pre-incubated with each streptavidin-fluorophore (all Biolegend) at a 1 µg/mL dilution before staining to allow tetramer formation, then combined and cells resuspend in the mixture and incubated for 30 min on ice, before staining for other surface markers. Samples were acquired on a BD LSRFortessa or BD Symphony A3 or A5. All cell sorting was performed on the BD FACSymphony S6. Data was analyzed using FlowJo 10.10.0.

### Recombinant protein production

All expression vectors for recombinant protein production were confirmed by Sanger (Azenta) or Oxford Nanopore sequencing (Plasmidsaurus). The construct encoding the RBD of SARS-CoV-2 (GenBank MN985325.1; S protein residues 319–539) was previously described (Barnes et al., 2020). All recombinant proteins were produced in Expi293F cells (Gibco, A14527), except 426c.WT.SOSIP which was produced in Expi293F GnTI-cells (Gibco, A39240). Transient transfections of expression plasmids were performed using Expifectamine 293 transfection kit (Gibco, A14525) as previously described (McGuire et al., 2014). In brief, 4-6 days post transfection, culture supernatant was harvested, centrifuged to pellet cells and sterilized by filtration for affinity chromatography. Trimeric Env proteins were purified by passing the supernatant through an agarose-bound *Galanthus nivalis* lectin (GNL) resin (Vector Laboratories) and subsequent size-exclusion chromatography (SEC). A Ni Sepharose 6 Fast Flow resin (Cytiva, 117531803) was used for purification of His-tagged proteins. Native gel electrophoresis identified peak fractions from size-exclusion chromatography. Fractions corresponding to monomeric Envelope trimers, Spike S6P protein trimers or RBD monomers were pooled and stored at −20 °C. Antibodies were purified over a protein G Sepharose 4Fast Flow resin (Cytiva 70611805) and buffer exchanged into PBS and stored at −80 °C.

For random biotinylation, proteins were biotinylated using the EZ-Link-Sulfo-NHS-LC-Biotinylation kit according to the manufacturer’s instructions (Thermo Fisher Scientific, 31497). Excess biotin was removed by diafiltration with 100 kDa cutoff. Biotinylated protein was stored at −20 °C or −80 °C.

Avi-tagged TM4 core gp120 or RBD were biotinylated using a BirA reaction according to the manufacturer’s instructions using a 5-fold molar excess of biotin (Avidity, EC6.3.4.15), buffer exchanged to PBS and stored at −80°C.

### ELISA

All ELISAs used Costar 96-well, half area, high binding, polystyrene assay plates (Corning, Cat.# 3960), which were coated with the indicated antigen (TM4, SARS-CoV-2 Wuhan-Hu-1 RBD or spike protein, all produced in house) or anti-isotype antibody (anti-IgM, Southern Biotech, 1020-01; anti-IgG3, Southern Biotech, 1100-01; anti-IgG1, Southern Biotech, 1070-01; anti-IgG2b, Southern Biotech, 1090-01; anti-IgG2c, Jackson Immunoresearch, 115-005-208; anti-IgA, Southern Biotech, 1040-01) at 2 – 5 µg/mL in PBS over night at 4°C. Plates were blocked with 5 % skimmed milk powder in PBS or 1 % BSA 0.1 mM EDTA 0.05 % Tween20 in PBS for 2 h at room temperature. 6 washes in PBS 0.05 % Tween 20 were performed after every subsequent step. Sera were diluted in PBS at 1:50 to 1:100 top dilution and 1:3 (naïve and after prime) or 1:4 (after boosts) or 1:5 (steady state serum antibody levels) serially diluted for an 8-point curve. Isotype antibody standards (Southern Biotech 5300-01B) were diluted to 10 µg/mL and diluted 1:5 in PBS for an 8-point curve. Secondary antibodies conjugated to horseradish peroxidase were used to detect bound antibodies mouse IgG (1:5000, Jackson ImmunoResearch Cat.# 115-035-071 or Southern Biotech Cat.# 1030-05) or mouse IgM, IgG3, IgG1, IgG2b, or IgA (Southern Biotech Cat.# 5300-05B) or IgG2c (Jackson ImmunoResearch 115-035-208) or total Ig (anti-kappa combined with anti-lambda light chain, Southern Biotech Cat.# 5300-05B). HRP substrate 3,3’,5,5’-tetramethylbenzidine substrate (ThermoFisher, Cat.# 34021) was used for development and the reaction was stopped adding an equal volume of 1 M H_2_SO_4_ (Sigma). Absorbance was read at 450 nm and 570 nm on a FLUOstar Omega (BMG Labtech). For analysis of steady state antibody concentration in serum, isotype standards were included on every plate and used to fit a sigmoidal 4-parameter logistic regression standard curve in GraphPad Prism 10 which was used to interpolate serum concentrations from dilutions in the exponential phase of the curve. For total anti-antigen responses, sigmoidal 4-parameter logistic regression was fitted to curves of every sample and used to calculate the half maximal binding titer (BT_50_). µg/mL values below detection were set to minimum detection level based on control sera to avoid plotting 0 on logarithmic plots. For samples where signal was too low for a curve fit BT_50_ was set to 10, which is 5x above the highest measured dilution.

### 10x genomics and single-cell libraries

Indicated mice were immunized with either Spike, RBD or HIV proteins. Cell suspensions were prepared as mentioned before from popliteal lymph nodes and samples were indexed using TotalSeqC cell surface antibodies (Biolegend). Live, Dump^−^ B cells (CD38^+^, GC B cells RBD^+^ and GCB RBD^−^), and PC were then isolated flow cytometry and loaded onto the Chromium Controller from 10x Genomics. scRNA-seq libraries were prepared using the Chromium Single Cell 5′ v.2 Reagent Kit (10x Genomics) according to the manufacturer’s protocol. Libraries were loaded onto an Illumina NovaSeq for single-cell gene expression (GEX), VDJ analysis and hashtags (HTO) library at the Rockefeller University Genomics Resource Center. Hashtag indexing from TotalSeqC antibodies was used to demultiplex the sequencing data and generate gene and barcode matrices, respectively.

All single-cell BCR libraries were mapped to the Cell Ranger VDJ GRCm38 reference using Cell Ranger multi v.8.1.0 (10x genomics). Contigs containing less than 50 reads and more than one heavy or light chain were removed. Antibodies heavy and light chains were paired using in-house scripts, and further analyzed using igpipeline v.3 (https://github.com/stratust/igpipeline/tree/igpipeline3) to define clonotypes, as previously described (Wang et al., 2023), using the mouse IMGT database as reference (Lefranc, 2011).

For the primary response dataset analyses (Fig. 1 H and Fig. 3), scRNA-seq and Hashtag-oligos unique molecular identifier quantification were performed with Cell Ranger multi v.8.0.1 using the Cell Ranger GEX reference mm10, and analyzed in R with Seurat v.5.1.0 (Hao et al., 2021). Cells were demultiplexed with MULTISeqDemux, and those classified as doublets or with mitochondrial content greater than 10 % and feature count less than 1,000 or greater than 6,500 were excluded. Cell cycle genes were regressed out. Sample batches were then merged, scaled and normalized with SCTransform. B cell (*Cd19, Ms4a1, Cd79a, Tnfrsf13c, Aicda, Lyz2, Cd63*), Follicular (*Cd38, Cd55, Bcl2, Maml2, Notch2, Gpr183, Sell, Ccr6*), and memory B cell (Binet et al., 2024) signatures were assigned using AddModuleScore. Cells with low B cell module score (≤ 25th percentile) or with high memory B cell score (≥ 75th percentile) were excluded. B cells expressing *Bcl6, Aicda, S1pr2* or *Fas* at high levels (≥ 75th percentile) were classified as GC B cells, whereas those with high follicular score (≥ 75th percentile) were classified as follicular B cells.

### Statistical analysis

Details of statistics including tests used, exact values and n numbers are indicated in figure legends and/or main text. Quantification and statistical analyses were performed in R (v4.4.0) with Rstudio server (2024.04.0 Build 735) and/or GraphPad Prism (Version 10.2.3), unless otherwise detailed in this methods section. Graphs generated using Prism and R were assembled into figures using Adobe Illustrator. Flow cytometry analysis was performed in FlowJo v.10.10.0 software (BD).

---

**Figure S1.**
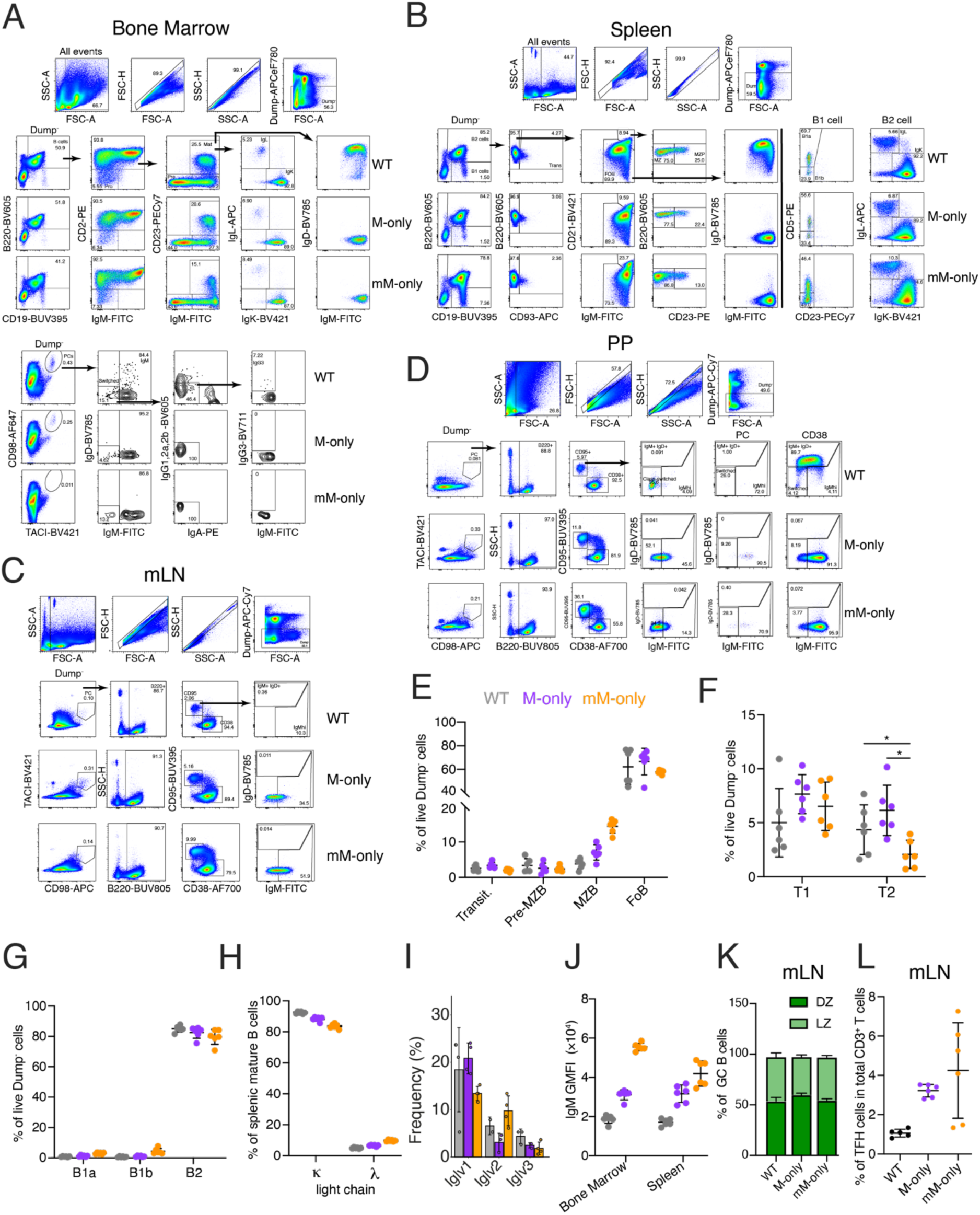
Analysis of M-only and mM-only mouse strains. Related to Fig. 1. (**A**) Flow cytometric gating strategy in the bone marrow. All stainings were cell surface and not intracellular stainings. (**B**) Flow cytometric gating strategy in the spleen. (**C**) Flow cytometric gating strategy in the mesenteric LN (mLN). (**D**) Flow cytometric gating strategy in the Peyer’s patches (PP). (**E**) Percentages of splenic, transitional (Transit), pre-marginal zone (Pre-MZB), marginal zone (MZB), and follicular (FoB) B cells among live Dump^−^ cells. (**F**) Percentages of transitional T1 and T2 B cells in live Dump^−^ cells. (**G**) Percentages of splenic B1a, B1b and B2 cells among live Dump^−^cells. (**H**) Percentages of splenic *κ* or *λ* light chain bearing mature B cells. (**I**) Bar graphs showing Iglv gene usage in follicular B cells. Top ten most frequent genes ranked by the mean frequency in mM-only mice are shown. (**J**) Geometric mean fluorescence intensity (GMFI) of IgM-FITC in the bone marrow mature B cells and splenic follicular B cells. (**K**) Proportions of DZ and LZ B cells in mesenteric lymph nodes at steady state. (**L**) Proportions of T follicular helper cells in mLN at steady state in total CD3^+^ T cells. Each dot represents a single mouse. Bars indicate mean ± SD.

**Figure S2.**
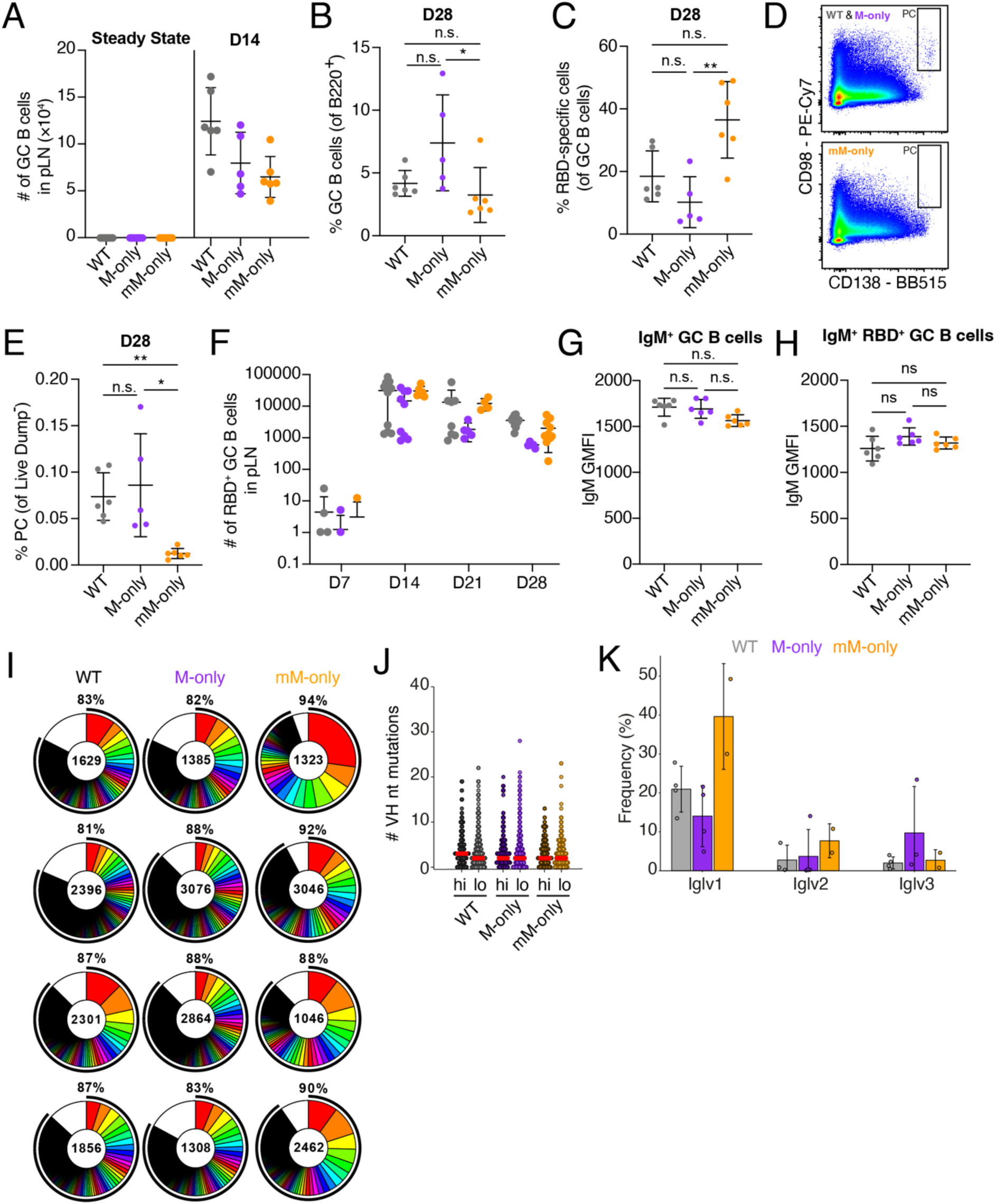
Prime immunization with RBD. Related to Fig. 2 and 3. (**A**) Absolute numbers of GC B cells in draining LN at steady state (left) and 14 days after RBD immunization (right). (**B**) Proportions of GC B cells in B220^+^ B cells at day 28 after immunization. (**C**) Proportions of RBD-specific cells in GC B cells at day 28 after immunization. (**D**) Flow cytometric gating strategy for PC. (**E**) Proportions of PC in live dump^−^ cells. (**F**) Numbers of RBD^+^ GC B cells at D7, D14, D21 and D28 after RBD immunization. (**G**) IgM geometric mean fluorescence intensity (GMFI) in CD95^+^ B220^+^ GC B cells. (**H**) IgM GMFI in RBD-specific CD95^+^ B220^+^ GC B cells. (**I**) Pie charts depicting the distribution of antibody sequences obtained 14 days after prime immunization from GC B cells from 4 mice/genotype. Inner circle numbers indicate the number of sequences analyzed for each mouse. White section indicates non-expanded sequences, colored or black pie slices are proportional to the number of clonally related sequences. The outlined black line indicates the percentages of cells in a clonal family. (**J**) Iglv gene usage frequency in genotypes. VH genes split by the mean frequency in mM-only are shown. Top 10 highest frequency (hi) and remaining low frequency VH genes (lo) are plotted. Each dots represents a VH gene. Bars indicate median. (**K**) Dot plot showing the number of VH nucleotide somatic hypermutations in sequences from top 10 most frequent V genes in mM-only mice from Fig. 5 F (Hi) compared with sequences from all remaining low-frequency V genes. Each dot indicates VH sequence from one cell. Bars indicate median. Each dot represents a single mouse. Bars indicate mean ± SD. *p<0.05, **p<0.01 in one-way ANOVA.

**Figure S3.**
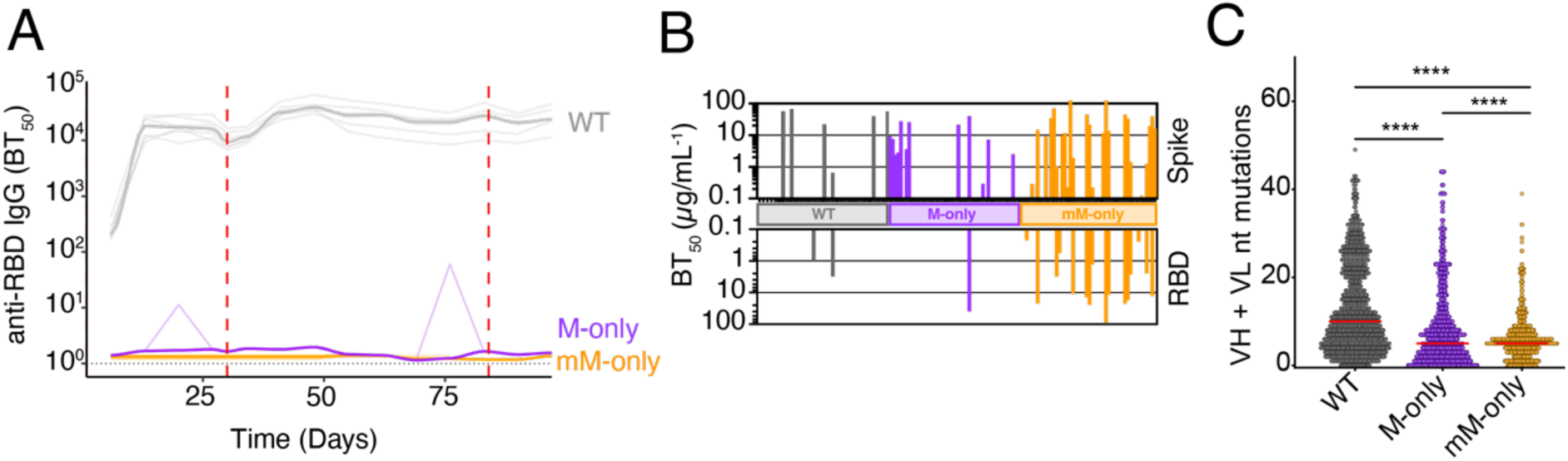
Analysis of anti-Spike prime boost responses. Related to Fig. 4. (**A**) ELISA quantification of anti-RBD IgG antibodies in the serum of mice from Fig. 4 measured weekly from day 6 to day 97. Thin lines represent single mice, bold lines represent group mean. (**B**) Bar graph of recombinant antibodies binding to Spike (top) or RBD (bottom) protein. Each bar represents one antibody. (**C**) Dot plot depicting somatic hypermutations in GC B cells on day 98 from Fig. 4. Each dot indicates one sequence. Bars indicate median. P-value of two-sided Mann-Whitney U test indicated. ****p<0.0001

**Figure S4.**
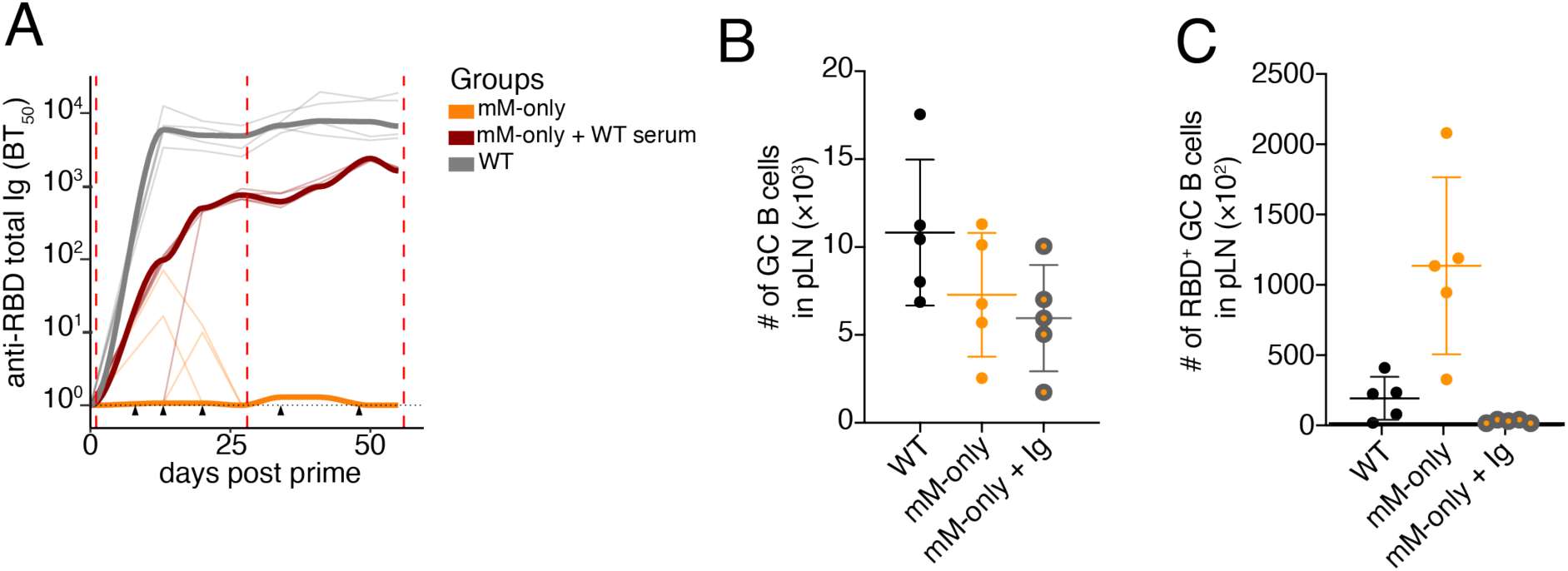
Analysis of serum transfer experiment. Related to Fig. 5. (**A**) ELISA quantification of anti-RBD total Ig antibodies in the serum measured weekly from day 0 to day 55. Red dashed lines indicate immunizations, black triangles indicate serum transfer. Thin lines indicate individual mice, bold lines indicate group mean. (**B**) Number of GC B cells in dLN. (**C**) Number of RBD-specific GC B cells in dLN. Each dot represents a single mouse, bars indicate mean ± SD.

**Figure S5.**
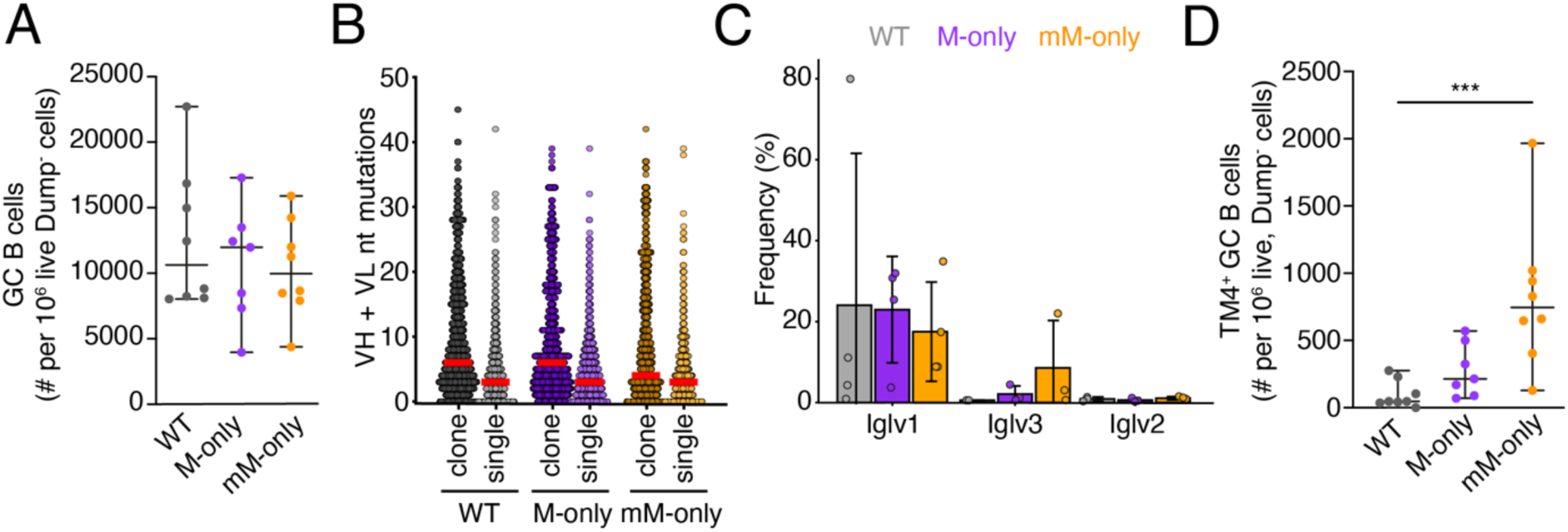
Analysis of anti-HIV-1 sequential immunization. Related to Fig. 5. (**A**) Number of GC B cells in draining lymph nodes after sequential immunizations on day 96. (**B**) Dot plot showing combined VH + VL gene nucleotide mutations among clonally expanded sequences compared with non-expanded sequences (singlets). Each dot indicates an antibody VH+VL pair. Bars indicate median. (**C**) Bar graphs showing Iglv gene usage in GC B cells. Top ten most frequent genes ranked by the mean frequency in mM-only mice are shown. (**D**) Number of TM4-binding GC B cells in sequentially immunized mice on day 96 in draining LNs. Each dot represents a single mouse, bars indicate mean ± SD unless otherwise indicated. ***p<0.001 non-parametric one-way ANOVA p-values are shown.

**Table S1. Recombinant antibody sequences and binding data.** Related to Fig. 4.

**Table S2. Flow Cytometric reagents.**

## Acknowledgements

We thank Kristie Gordon and Jean-Philip Truman for assistance with cell sorting, Thomas Eisenreich for help with mouse colony management and technical help; Masa Jankovic and Tacio A. Waldetario for laboratory support; and all members of the Nussenzweig laboratory and HIVRAD P01 (2P01AI100148) collaborators for helpful discussions.

## Author contributions

D.S.B., H.H. and M.C.N conceived the study. D.S.B and H.H. generated the novel mouse strains described in this manuscript. D.S.B., L.B. and H.H. designed and performed the experiments and interpreted data. G.S.S.S. performed bioinformatic analysis. C.R., L.D., M.A.E., D.G., G.L.D.R., and C.U. assisted and performed experiments. K.H.Y. assisted with mouse experiments and husbandry; B.H. and A.G. produced recombinant proteins and antibodies. P.A. and L.S. provided HIV reagents. M.C.N. supervised the study, designed experiments and interpreted data. D.S.B., L.B., H.H. and M.C.N. wrote the manuscript. All authors edited the manuscript.

## Data and materials availability

All data needed to evaluate the conclusions in the paper are available in the main text and supplementary materials. Reagents, including mouse strains, are available from H.H. and M.C.N. under a material transfer agreement with The Rockefeller University under reasonable request.

## License information

This article is subject to HHMI’s Open Access to Publications policy. HHMI lab heads have previously granted a nonexclusive CC BY 4.0 license to the public and a sublicensable license to HHMI in their research articles. Pursuant to those licenses, the Author Accepted Manuscript (AAM) of this article can be made freely available under a CC BY 4.0 license immediately upon publication.

